# Reprogramming Cas9 PAM Recognition for Allele-Specific Editing

**DOI:** 10.64898/2026.08.10.744034

**Authors:** Julia A. Tartaglia, Vivian Nguyen, John J. Desmarais, Rachel F. Weissman, Brittney W. Thornton, Marena I. Trinidad, Kevin Briseno, Taylor R. Hudson, Carmelle Catamura, Liana F. Lareau, Fyodor Urnov, Jennifer A. Doudna, David F. Savage

## Abstract

The therapeutic potential of CRISPR–Cas9 genome editing is fundamentally constrained by the requirement for specific short DNA sequences (PAMs) flanking the target site, limiting access to many clinically relevant genomic loci. This stringent PAM requirement is particularly problematic in applications which require precise positioning, such as base editing and allele-specific editing. Although PAM-relaxed variants have expanded the targetable genome, they incur trade-offs in on-target activity, off-target editing, and cleavage kinetics. This highlights an unmet need for variants that are re-targeted to alternative PAMs in order to maintain the specificity and enzymatic performance inherent to stringent dinucleotide PAM recognition. To overcome these limitations, we developed a yeast selection platform to engineering SpCas9 variants with re-specified PAM recognition. Using a clinically relevant Huntington’s disease gene (*HTT*) SNP as a proof-of-concept target, we engineered variants with reciprocal NGC and NGT PAM selectivity, as a step toward allele-specific editing in a large percentage of Huntington’s disease patients. These yeast-selected SpCas9 variants retained their modified activity across multiple endogenous HEK293T loci, demonstrating that this specificity is robust across diverse genomic contexts. The variants surpassed PAM-broadened variants on their respective on-target PAM while displaying broad loss of activity across alternative PAMs, effectively re-specifying PAM recognition toward a single dinucleotide sequence. Retargeted variants recovered on-target cleavage kinetics approaching that of wild-type SpCas9, even under competing substrate conditions, demonstrating that PAM re-specification can simultaneously restore catalytic efficiency and improve specificity. Beyond NGC and NGT, we leveraged our high-throughput platform to engineer Cas9 with re-specified activity across multiple additional non-canonical PAMs in yeast, further demonstrating its utility as a general and programmable framework for expanding the therapeutic reach of precision genome editing.

## Introduction

RNA-guided CRISPR-Cas endonucleases enable precise and programmable changes to the genome^1^. This system offers immense therapeutic promise towards treating numerous genetic diseases. CRISPR-Cas9’s ability to target a specific location within a genome is conferred by a guide RNA (gRNA), which binds Cas9, forming a ribonucleoprotein complex (RNP)^1–4^. Proper targeting requires both a 20-nucleotide gRNA spacer base-pairing with complementary DNA and the presence of a protospacer adjacent motif (PAM) immediately following the target sequence (protospacer) for Cas9 binding and cleavage (Figure 1A)^1,5–7^. This PAM dependence evolved as a fundamental mechanism for self–non-self discrimination, enabling selective targeting of invading nucleic acids while avoiding cleavage of the host genome. *Streptococcus pyogenes* Cas9 (SpCas9), the most widely used genome editing tool, recognizes NGG (N = A, T, C, or G) through specific and non-specific electrostatic interactions with its PAM-interacting domain^8–11^, imposing accessibility constraints that restrict the editable sequence space and limit therapeutic applications such as base editing and allele-specific editing.

**Figure 1:**
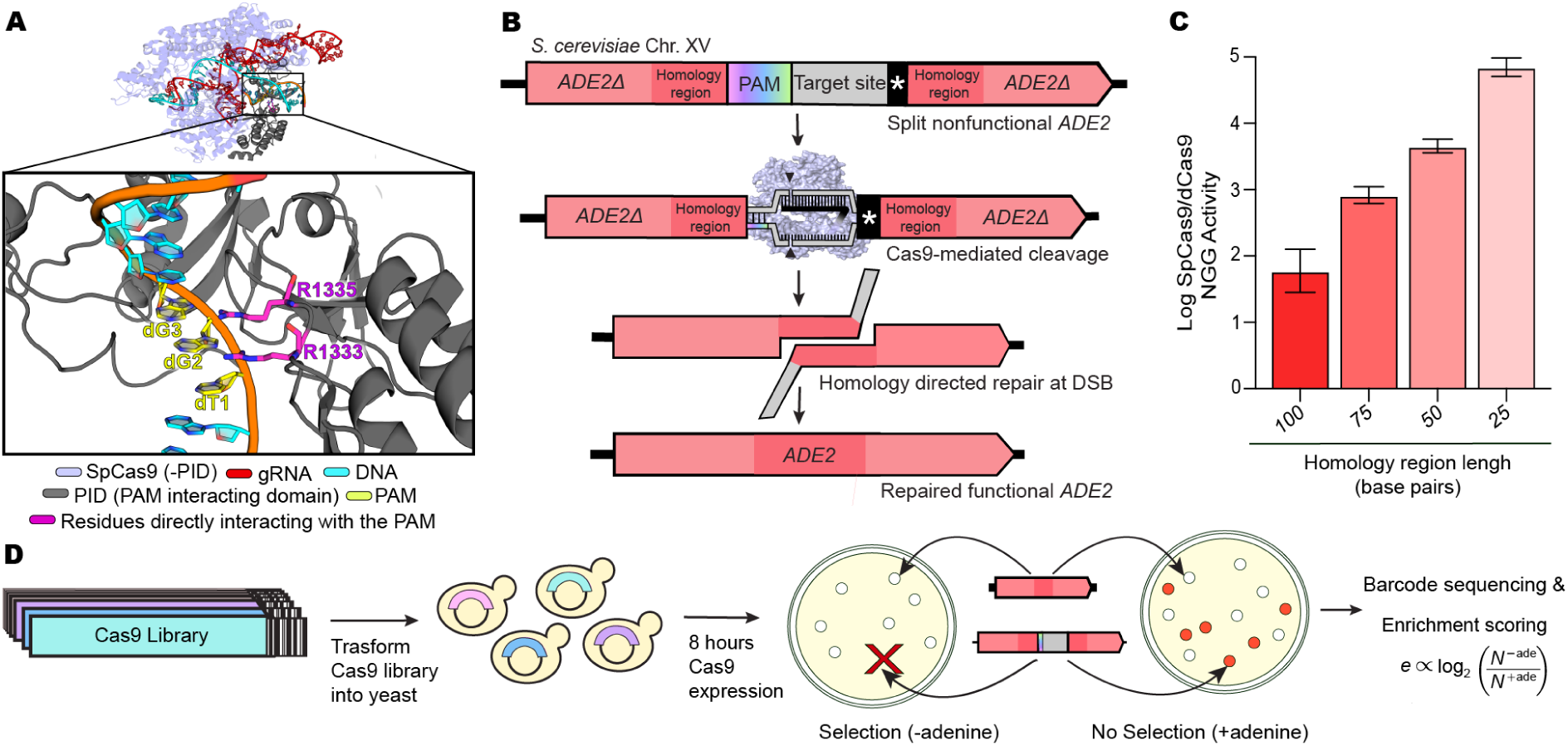
Selection system optimization and workflow. **(A)** Domain organization of SpCas9. Ribbon representation of SpCas9-sgRNA-DNA ternary complex. sgRNA in red; DNA in cyan; PAM-interacting domain in grey and the rest of Cas9 in purple. Zoomed view highlights the PAM-binding site in yellow with direct PAM interacting residues highlighted in pink. Structure from PDB ID 4UN3. **(B)** Design of yeast reporter strains. A cassette containing any PAM or spacer of interest is inserted into the *ADE2* gene along with a stop codon (*) and flanking homology regions. Colony survival under selective (–adenine) conditions indicates successful PAM recognition, cleavage, and homology-directed repair. **(C)** Optimization of homology arm length in reporter strains to maximize dynamic range between functional and non-functional variants. Wild-type or dead SpCas9 was transformed into strains carrying different homology arm lengths and a NGG PAM, stably expressing an on-target sgRNA. Editing efficiency was determined from colony counts on selective versus non-selective plates (Figure S1C). Percentages (± s.e.m.) from n = 3. **(D)** Pooled yeast screening workflow for differential variant activity at a specific PAM. Cas9 variant plasmid libraries, each with a unique 20-bp C-terminal barcode, were transformed into *ADE2* reporter strains stably expressing a gRNA. Post-expression, libraries were plated under selective (–adenine) and non-selective (+adenine) conditions. Barcode sequencing and log-ratio analysis of selective/non-selective abundance determined relative enrichment for each variant.

Previous work has expanded SpCas9’s targeting range by broadening PAM specificity^12–14^, yielding novel variants such as SpRY Cas9 that target PAMs with a purine, and to a lesser extent a pyrimidine, at the second PAM position^14^. However, reducing the PAM requirement to fewer than two specified nucleotides results in lower on-target activity, slower kinetics, and higher off-target activity, limiting therapeutic potential^15–17^. A number of non-NGG specific PAM targeting variants have been identified through mining of natural Cas9 orthologs as well as directed evolution of SpCas9, but such work has largely yielded longer and purine-rich PAMs^18–22^. Machine learning approaches have also been employed to investigate key residues within the PAM-interacting domain that preserve PAM specificity while detargeting recognition away from NGG^23^. Despite these advances, the ability of CRISPR–Cas9 systems to target the majority of the genome with high activity and specificity remains to be established.

A particularly compelling application for PAM-diversified Cas9 variants would be the use of allele-specific genome editing to selectively modify a disease-causing allele while preserving its healthy counterpart. This distinction is critical in dominant disorders, where indiscriminate editing of both alleles would ablate residual wild-type function and potentially exacerbate disease^24^. Broad, degenerate PAM-targeting Cas9 variants lack the specificity required to discriminate between alleles that differ at their PAM sequence, whereas v^5,8,25,26^ ariants with strict PAM-dependent recognition can selectively engage the disease allele while leaving the healthy allele unedited. The most stringent allelic discrimination can be achieved when the disease-causing mutation resides at the PAM sequence, yet without PAM-diversified Cas9 variants this strategy is confined to the small subset of allelic mutations that coincide with an NGG PAM^27–29^.

As a proof of concept for evolving allele-specific PAM-targeting variants in a therapeutically relevant context, we selected Huntington’s disease, a condition that is largely inaccessible to current genome-editing strategies. Huntington’s disease is a fatal dominant genetic disorder phenotypically characterized by progressive breakdown of neuronal cells and genotypically characterized by the expansion of tandem trinucleotide (CAG) repeats to greater than 36 copies on the diseased allele of the Huntingtin (*HTT*) gene^30^. While SpCas9 20 bp target sequence cannot differentiate between a normal and diseased number of repeats, heterozygous single nucleotide polymorphisms (SNPs) in the *HTT* gene provide an alternative recognition site for specific targeted knockout of the diseased Huntington’s allele^31,32^. Previous studies have exploited SpCas9’s NGG preference for allele-specific *HTT* targeting, achieving selective knockout and reduction of Huntington’s disease phenotypes, but NGG-containing SNPs at the genomic positions required for such targeting are rare, limiting broad therapeutic applicability^31,32^. Rarer still are SNPs placing variation within the PAM itself — a critical consideration, as variation at the second and third PAM positions confers the strongest allelic discrimination. Here, we tested the ability to discriminate existing *HTT* SNPs abundantly found in the human population that require high and specific Cas9 activity on sites with either NGC or NGT PAMs (Lareau lab, manuscript in preparation). This is currently difficult with existing Cas9 technology, which largely target NRR PAMs. To this end, we describe here the engineering and application of novel Cas9 variants re-targeted to only NGC or NGT activity, with discrimination against other PAMs, and thus suitable for allele-specific editing.

To evolve SpCas9 variants with re-specified PAM recognition capable of therapeutically relevant allele-specific editing, we designed and optimized a selection system in *Saccharomyces cerevisiae*^33–35^, enabling the screening of large libraries of Cas9 variants for selective targeting of any PAM sequence and protospacer of interest. Libraries were generated through random mutagenesis of the entire PAM-interacting domain of SpRY Cas9, with increased mutational frequencies at residues previously shown to influence PAM activity^14,18^. The PAMless nature of SpRY Cas9 provides substantial evolutionary plasticity, potentially enabling both enhanced targeting of weakly recognized sequences and loss of activity at previously targeted sites^14,36–38^. These libraries were selected for allele-specific targeting of the *HTT* site, allowing the identification of Cas9 variants with enhanced activity on NGC while minimizing activity on NGT, and vice versa. Comprehensive dinucleotide PAM profiling and biochemical analysis of the top allele-specific variants revealed the general de-targeting of all PAMs except the on-target PAM, and improved enzyme kinetics on the targeted PAM. In human cells, variants identified from the selection showed strong PAM selectivity for their on-target over their off-target PAM on numerous endogenous sites, with increased editing at desired editing sites and lower genome-wide off-target activity than SpRY Cas9. These results highlight the capacity of our screening approach to evolve novel Cas9 variants with differential PAM selectivity in a therapeutically relevant, allele-specific disease context.

## Results

### Engineering of NGC > NGT and NGT>NGT PAM-targeting Cas9 libraries

To engineer variants capable of distinguishing NGC from NGT PAMs, we selected SpRY Cas9 as our mutational scaffold, as its broad PAM compatibility provides a foundation for fine-tuning specificity in either direction without requiring evolution of entirely new activity. Libraries were generated by first mutating residues 1333 and 1335, which contact the second and third PAM nucleotides, respectively, biased toward amino acids favoring the desired PAM bases^8,39–46^. Given that mutations at direct PAM-contact residues alone are insufficient to fully shift PAM preference while retaining robust activity, we additionally mutated residues with known influence on PAM recognition (1135, 1136, 1218, 1219, and 1337) at higher frequencies. This rationally designed library was then expanded via error-prone PCR to incorporate second-and third-shell mutations throughout the PAM-interacting domain, balancing targeted mutagenesis of known contact residues with broader sampling to capture compensatory combinations and unknown residues influencing domain stability and active-site conformational dynamics. Critically, introducing multiple mutations simultaneously enabled access to epistatic effects and novel PAM-selective combinations inaccessible to deep mutational scanning or sequential site-directed mutagenesis approaches. Each variant was uniquely barcoded with a random 20-bp sequence for high-depth short-read sequencing, with Cas9 mutations mapped to barcodes via long-read PacBio sequencing (see Methods).

### Optimization and validation of a yeast-based endonuclease selection system

To assess PAM recognition and cleavage activity of Cas9 variants, we adapted a *Saccharomyces cerevisiae* reporter strain previously used to engineer Cas9 variants with reduced off-target activity^33^. The endogenous essential *ADE2* gene was split by insertion of a cassette containing a defined target spacer and PAM sequence, a stop codon, and flanking homology arms. This design created a selection system in which cell survival and proliferation under adenine-deficient conditions strictly depends on Cas9-mediated cleavage and subsequent homology-directed repair at the reporter locus (Figure 1B). Systematic shortening of the homology arms to 25 bp markedly reduced background recombination, thereby optimizing selection stringency and sensitivity at lower activities (Figures 1C and S1C). To validate the system’s performance, we evaluated wild-type SpCas9 alongside HypaCas9, HypaCas9+928A, SpRY, and dCas9 — variants with well-characterized differences in activity — in both pooled (Figure 1D) and individual formats^1,14,47,48^. The assay robustly resolved differential editing efficiencies among these variants, demonstrating its sensitivity and scalability for quantitative assessment of Cas9 activity (Figures S1A-C).

### Selection of NGC and NGT PAM-targeting variants in yeast

To select for Cas9 variants with PAM specificities for the proof-of-concept *HTT* alleles, we generated two *ADE2* yeast reporter strains containing the aforementioned SNPs resulting in either an NGC or NGT PAM (Figure 2A). Plasmids encoding the guide RNA and pooled NGC/NGT Cas9 variant library, representing a diversity on the order of 10^5^ variants, were transformed into reporter strains with NGC or NGT PAMs. Cas9 expression was then induced for 8 hours, and then transformed cells were plated on selective (–adenine) and nonselective (+adenine) media. Barcodes linked to individual Cas9 variants were amplified from yeast recovered from each condition and quantified by short read sequencing. For each variant, relative enrichment on NGC and NGT PAMs was calculated as the ratio of variant abundance under selective versus nonselective conditions, averaged across independent replicates (Figures 2B and S2A). Variants exhibiting the greatest differential enrichment between the NGC- and NGT-targeting reporter strains were isolated from the library using barcode-specific primers and individually evaluated for editing efficiency on NGC and NGT PAMs in yeast (Figures S2B-D). The top NGC-over-NGT–selective variants displayed up to 450-fold NGC/NGT activity, corresponding to a 380-fold increase relative to SpRY, whereas the top NGT-over-NGC–selective variants achieved up to 180-fold NGT/NGC activity, a 220-fold improvement over SpRY (Figure 2C and 2D).

**Figure 2:**
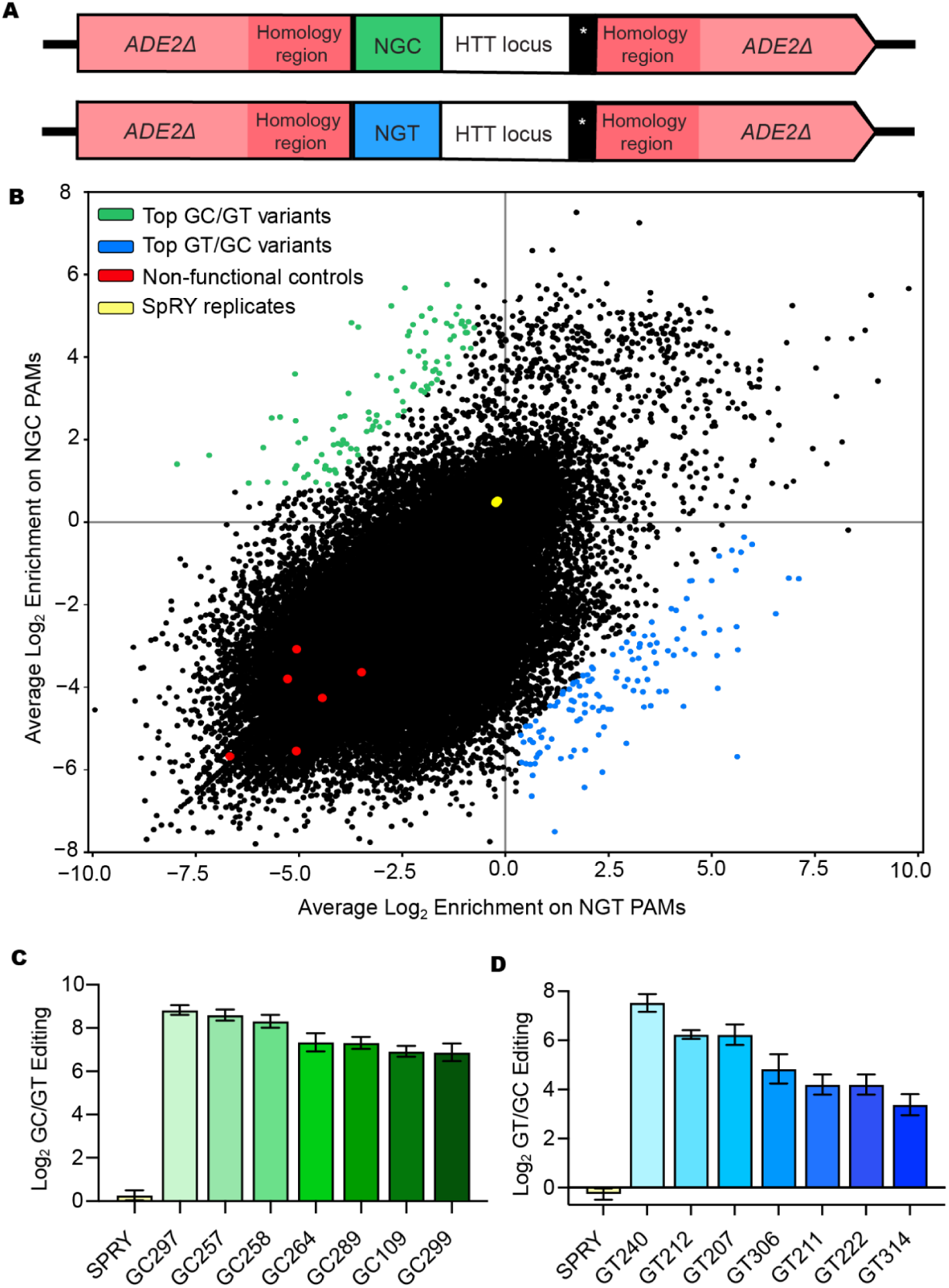
High-throughput screening identifies allele specific PAM-targeting Cas9 variants. **(A)** Design of the yeast reporter constructs used to assess Cas9 NGC and NGT PAM activity on a protospacer matching the *HTT* locus. A cassette containing this protospacer, a stop codon, and flanking homology regions was inserted into the *ADE2* gene together with either an NGC (green, top) or NGT (blue, bottom) PAM. **(B)** Variant libraries of SpRY Cas9, in which the PAM-interacting domain was mutagenized via combined random and rational design, were evaluated in NGC and NGT PAM reporter strains. The barcode unique to each Cas9 variant, located at the C-terminus, was PCR-amplified and sequenced using next-generation sequencing (NGS). For each variant, log_2_ enrichment was computed from barcode abundances in selective relative to nonselective conditions across two independent replicates. Each dot represents the mean relative enrichment (log_2_ > 0: enriched; log_2_ < 0: depleted). Non-functional controls in yellow; SPRY replicates in red; top NGC/NGT variants in green; top NGT/NGC variants in blue. **(C)** NGC/NGT PAM selectivity ratios for top NGC-targeting variants and **(D)** NGT/NGC PAM selectivity ratios for top NGT-targeting variants identified from the high-throughput screen, with SpRY shown for comparison. Selectivity ratios were calculated as the log_2_ ratio of mean editing on NGC and NGT PAMs. Variants were individually evaluated in *ADE2* reporter strains containing either NGC or NGT PAMs, and activity (mean ± s.e.m.) was determined from colony reversion rates from n = 3 replicates (Supplementary Fig. 2b–d).

### Evaluation of complete dinucleotide PAM-targeting profiles in yeast

To comprehensively define the PAM targeting capabilities of our top variants, we generated 16 *ADE2* yeast reporter strains representing all possible dinucleotide PAMs with an identical target sequence. Each strain encoded a unique PAM-identifying barcode downstream of the *ADE2* coding sequence for unambiguous locus identification following Cas9-mediated editing (Figure 3A). Strains were pooled, and top variants were individually transformed, transiently expressed, and plated under selective and nonselective conditions. Relative enrichment values for each dinucleotide PAM were calculated from barcode sequencing as the ratio of strain abundance under selective versus nonselective conditions (Figures 3B-D). Consistent with initial screening results, strong NGC/NGT and NGT/NGC PAM preferences were observed for all top variants. GC297, our leading NGC/NGT variant, exhibited a 25-fold NGC/NGT activity ratio, while GT240, our leading NGT/NGC variant, displayed an 88-fold NGT/NGC ratio, approaching the NGG/NGA selectivity of wild-type SpCas9. Interestingly, variants evolved for selective recognition of one PAM over one other and exhibited markedly reduced activity across the remaining 14 dinucleotide PAMs — with on-target PAM/NNN enrichment ratios approaching those of wild-type SpCas9 at NGG — indicating a general shift toward increased specificity and reduced PAM degeneracy.

**Figure 3:**
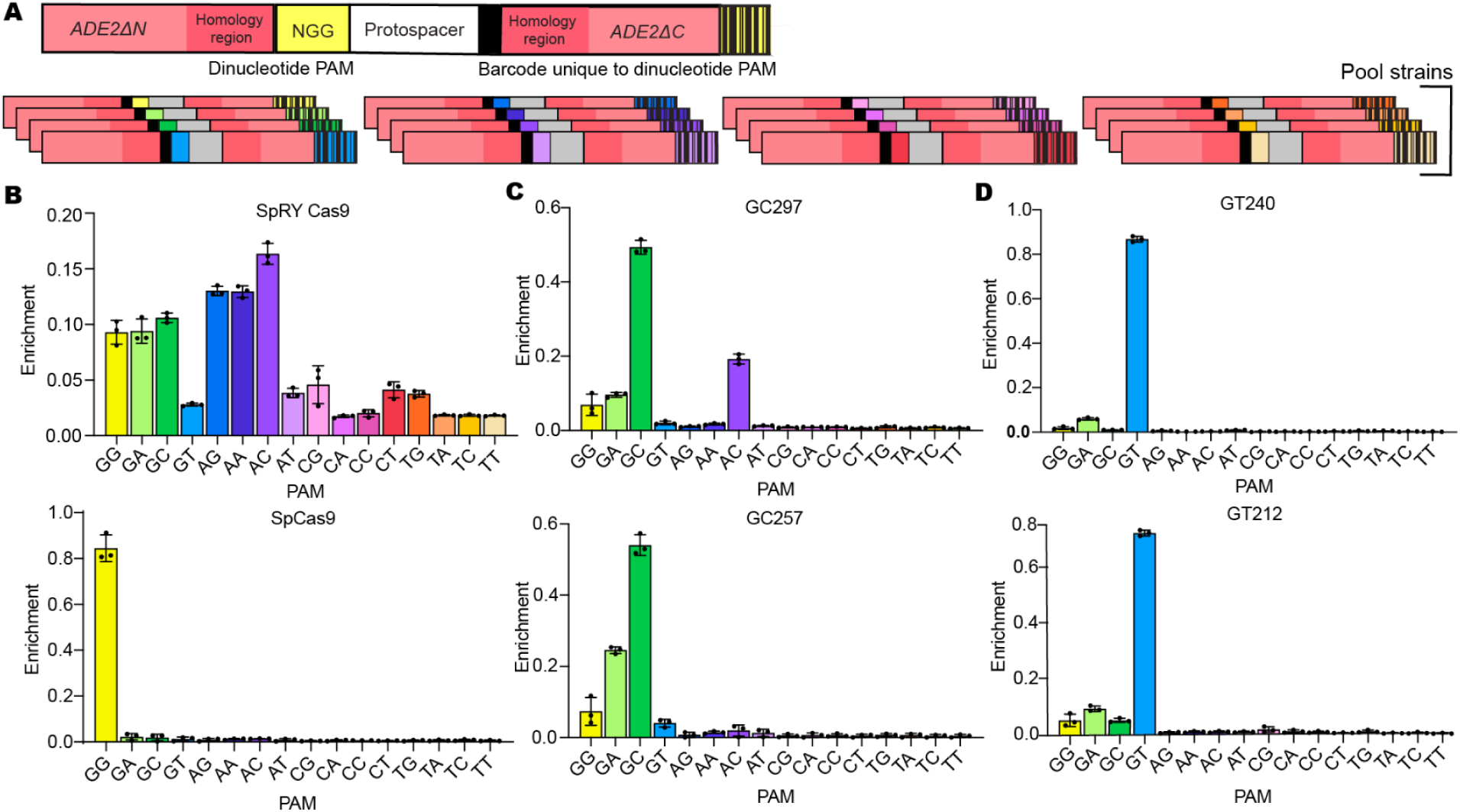
Selection for one PAM over another results in general detargeting of other dinucleotide PAMs. **(A)** Schematic of the 16 *ADE2* yeast reporter strains representing all possible dinucleotide PAMs with an identical target sequence, each carrying a unique PAM-identifying barcode at the C-terminus of the *ADE2* reporter gene for unambiguous locus identification following Cas9-mediated editing. **(B–D)** All barcoded PAM strains, stably expressing gRNAs, were pooled, and variants of interest were individually transformed. Post-selection, the *ADE2* C-terminal barcode region was sequenced, and the log-ratio of barcode abundance in selective versus non-selective conditions was used to calculate variant enrichment. Enrichment values are normalized such that the sum across all PAMs equals 1. Complete dinucleotide PAM profiles for: **(B)** control variants (SpRY Cas9, SpCas9) **(C)** top NGC/NGT-targeting variants (GC297, GC257), and **(D)** top NGT/NGC-targeting variants (GT212, GT240) in our yeast selection platform. Enrichment values (± s.e.m.) are from n = 3 independent selections.

### Extending the yeast selection platform to diverse dinucleotide PAMs

Having constructed selection strains targeting all dinucleotide PAMs, we asked whether our platform could be extended to engineer variants with re-specified recognition of additional noncanonical dinucleotide PAMs. As two-base PAMs have been shown to provide optimal trade-off between efficient search kinetics and broad genomic targetability^15,17^ we constructed libraries targeting dinucleotide PAMs not recognized well by both wild-type SpCas9 and recently engineered variants. Pyrimidine-rich PAMs are notoriously difficult to target with specificity, given their substantial divergence from the purine-rich PAMs that SpCas9 naturally evolved to recognize^12–14,18,49–51^. Methods capable of evolving variants that efficiently target these diverse sequences without incurring the costs of PAM relaxation would therefore be of great interest to the field. PAM-targeting libraries were generated by increasing mutational frequencies at residues known to influence PAM activity, with positions 1333 and 1335 biased toward amino acids reported to interact with the corresponding second and third PAM bases, respectively, followed by random mutagenesis across the SpRY PAM-interacting domain to expand sequence diversity. These libraries were subsequently screened in our *ADE2*-based yeast reporter system to determine enrichment of each variant on its target noncanonical PAM relative to a single-nucleotide-differing PAM from which it was de-targeted.

Top-performing variants showing high on-target and low off-PAM activity in the primary screen were cloned individually and re-validated against both their target and de-targeted PAMs. Among these, CG544 and AC495 edited NCG and NAC PAMs, respectively, with activity approaching that of wild-type SpCas9 at canonical NGG PAMs, demonstrating that our selection platform can isolate highly active variants for noncanonical PAM targeting (Figures S3A and S3C). Additional variants showed significantly greater activity on NCC, NTG, and NTC PAMs relative to SpRY, underscoring the platform’s versatility in generating retargeted variants for pyrimidine-rich PAMs and PAMs with T at the second position. (Figures S3A and S3C). Beyond their enhanced on-target activity, all variants showed markedly reduced activity on the single-nucleotide PAM from which they were de-targeted, indicating that the selection system can simultaneously optimize activity and specificity at desired PAMs (Figures S3B and S3C). Together, these results highlight our platform as a broadly applicable approach for engineering re-specified PAM recognition across diverse sequence contexts in yeast.

### Endogenous NGC and NGT PAM editing in HEK293T cells

To further characterize the performance of these variants, we selected the lead NGC- and NGT-specific variants to assess their re-specified activity in human cells. To evaluate their editing efficiency across diverse genomic contexts, we selected four endogenous human genomic loci requiring NGC PAM recognition and four requiring NGT PAM recognition. Plasmids encoding either a top-performing or a control Cas9 variant (either SpRY or dCas9) were co-transfected with a plasmid expressing a guide RNA targeting a single endogenous locus. Five days post-transfection, insertion–deletion (indel) frequencies were quantified by PCR amplification and sequencing of the targeted genomic regions (Figures 4A-C). The top NGC-over-NGT variants, GC297 and GC257, exhibited significantly higher editing efficiencies across the four NGC PAM–targeting loci, with average indel frequencies of 13%, compared with editing at NGT PAM–targeting loci, which averaged 0.3% and 0.5% respectively. Conversely, the top NGT-over-NGC variants, GT240 and GT212, displayed significantly higher editing across the four NGT PAM–targeting loci, averaging 22% and 26%, relative to editing at NGC PAM–targeting sites, which averaged 2% and 3% respectively. Notably, all engineered variants showed markedly higher on-target editing at the PAM they were specifically selected to recognize at all sites evaluated compared with SpRY. Of the two top NGC variants, GC297 had both the greatest overall NGC editing and the lowest NGT editing. Among the two top NGT variants, GT212 showed the highest NGT editing, whereas GT240 had lower NGC editing and a higher NGT/NGC editing ratio.

**Figure 4:**
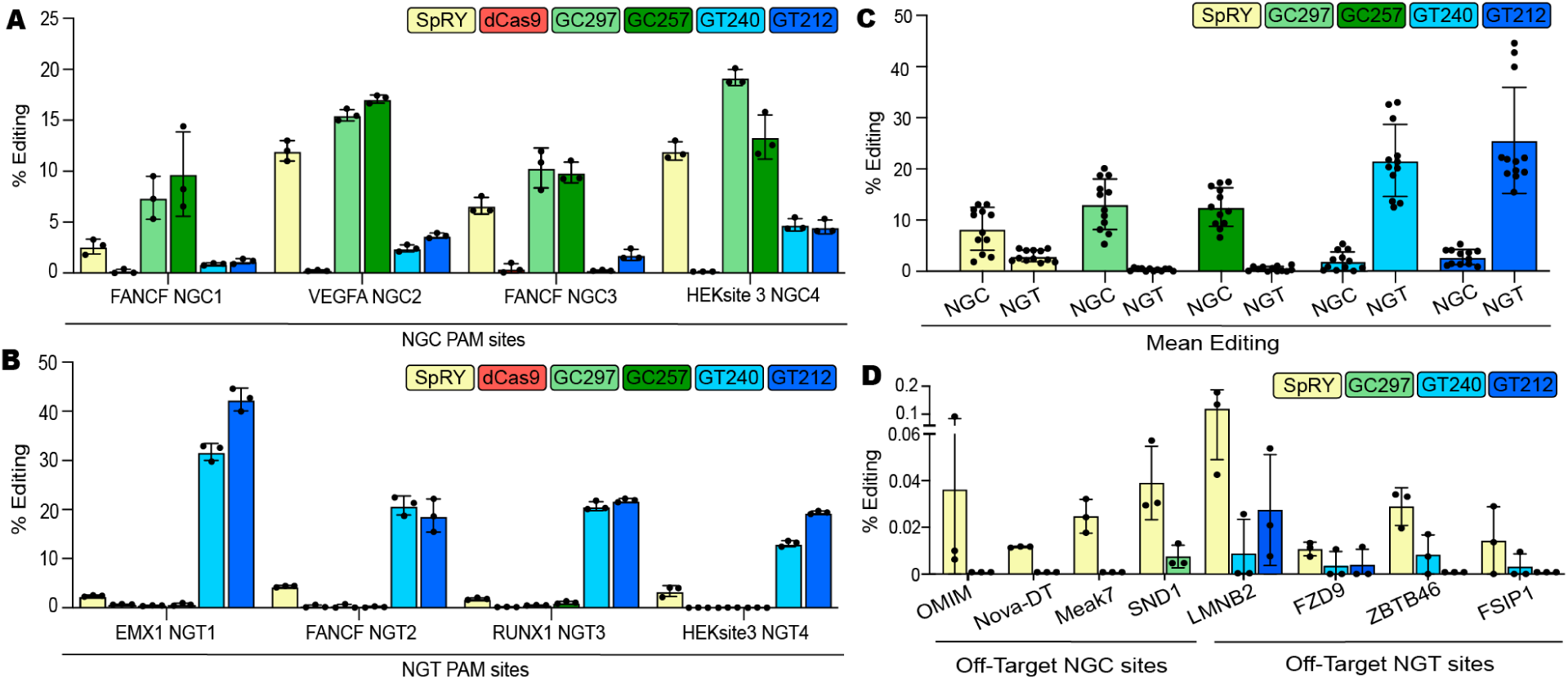
Top variants identified in yeast are active and PAM selective in human HEK293T cells. (A–B) Top two NGC/NGT and NGT/NGC PAM-targeting variants, along with control variants SPRY and dCas9, were evaluated for editing efficiency at four endogenous NGC PAM loci and four NGT PAM loci in HEK293T cells. Indel frequency was determined by targeted NGS of PCR-amplified genomic regions and analyzed with CRISPResso2. Data are mean ± s.e.m. from n = 3 biological replicates. **(C)** Editing profiles of the top NGC/NGT and NGT/NGC variants across multiple endogenous loci with NGC or NGT PAMs. **(D)** Top off-target sites in HEK293T cells for guides targeting two NGC PAM loci (for SPRY, dCas9 and GC297) and NGT PAM loci (for SPRY, dCas9, GT212, and GT240) were selected from sites predicted by Cas-OFFinder. Sites were evaluated for percent editing by targeted NGS of PCR-amplified genomic regions and analyzed with CRISPResso2. Data are presented as mean ± s.e.m. from n = 3 biological replicates.

### Editing at top predicted off-target sites in HEK293T cells

Minimal off-target activity is essential for safe therapeutic application. We therefore further assessed editing at four predicted off-target sites for our NGC and NGT on-target protospacer sequences (Figure 4D). SpRY exhibited detectable activity at all four predicted off-target sites for both NGC and NGT on-target guides. In contrast, GC297 showed off-target activity at only one NGC PAM off-target site, and at levels significantly below SpRY. For NGT guides, GT240 displayed reduced off-target activity relative to SpRY across all four sites, while GT212 showed detectable activity at only two sites, both below SpRY levels. The consistently lower off-target profiles of our variants compared to SpRY is encouraging for their therapeutic applicability.

### Genome wide off-target evaluation

To evaluate genome-wide off-target activity, we performed CHANGE-seq^52^ a biochemical assay in which the GC297, GT212, GT240, and SpRY nucleases complexed with guide RNAs targeting loci adjacent to their respective on-target PAMs, are incubated with purified human genomic DNA and cleavage sites identified. All engineered variants exhibited marked improvements in on-target to off-target editing ratios compared with SpRY, with increases of up to 54-fold for GT240, 42-fold for GT212, and 3-fold for GC297 (Figure S4A), accompanied by a substantial reduction in the total number of detected unique off-target sites. Notably, this reduction in unique off-target sites held even for the variant and sgRNA with the smallest improvement in on-target to off-target editing ratio relative to SpRY (Figure S4B).

In addition to their substantially reduced off-target activity relative to SpRY, a comprehensive analysis of each variant’s off-target PAM preferences further revealed pronounced re-targeting toward the PAM for which it was evolved, alongside reduced activity at the de-targeted PAM (Figures S4C and S4D). Among NGT-targeting variants, GT212 showed higher overall NGT PAM selectivity while GT240 displayed greater NGT/NGC PAM selectivity. GC297 exhibited particularly strong NGC/NGT PAM selectivity, with 18-fold NGC/NGT enrichment for GCgRNA1 and no detectable NGT off-target sites for GCgRNA2. This re-targeting was even more pronounced when restricting the analysis to the most frequently cleaved off-target sites (Figure S4E).

### Kinetic characterization of top Cas9 variants

Previous work measuring R-loop–limited DNA cleavage kinetics has shown that the highly promiscuous editor SpRY exhibits substantially slower on-target cleavage rates compared with NGG-specific SpCas9^15,17^. This reduction in activity is further exacerbated in the presence of competing substrates^15^. To determine whether re-specifying PAM recognition restores rapid on-target cleavage, we measured the R-loop–limited DNA cleavage kinetics of our top engineered variants.

Equally active enzyme fractions of purified Cas9 variants were complexed with guide RNA and incubated with MgCl_2_, and *in vitro* cleavage reactions were initiated by addition of a 5′ FAM-labeled 55-bp DNA substrate containing a perfectly matched protospacer and the PAM of interest (Figures S5A and S5B). Time-course analyses revealed both variant- and PAM-dependent differences in cleavage kinetics, from which rate constants (*k*_obs_) were determined (Figure 5A-D and S5B). Kinetic analyses revealed that GC297, the top NGC-over-NGT variant, cleaved its on-target NGC substrate 36-fold faster than SpRY, with a 165-fold difference between its on- and off-target PAM cleavage rates and an 87-fold higher NGC/NGT cleavage rate ratio relative to SpRY (Figures 5A and 5B). The top NGT-over-NGC variants, GT240 and GT212, exhibited 36- and 55-fold faster cleavage than SpRY on NGT substrates, respectively, with NGT/NGC cleavage rate ratios of 77- and 41-fold — representing 144- and 77-fold higher NGT/NGC cleavage rate ratios compared to SpRY (Figure 5A and 5C-D). Strikingly, on-target cleavage rates for all top variants approached those of wild-type SpCas9 on NGG (Figure 5A-E), demonstrating that re-targeting of SpRY toward a single dinucleotide PAM can restore rapid enzymatic kinetics while achieving high PAM selectivity.

**Figure 5.**
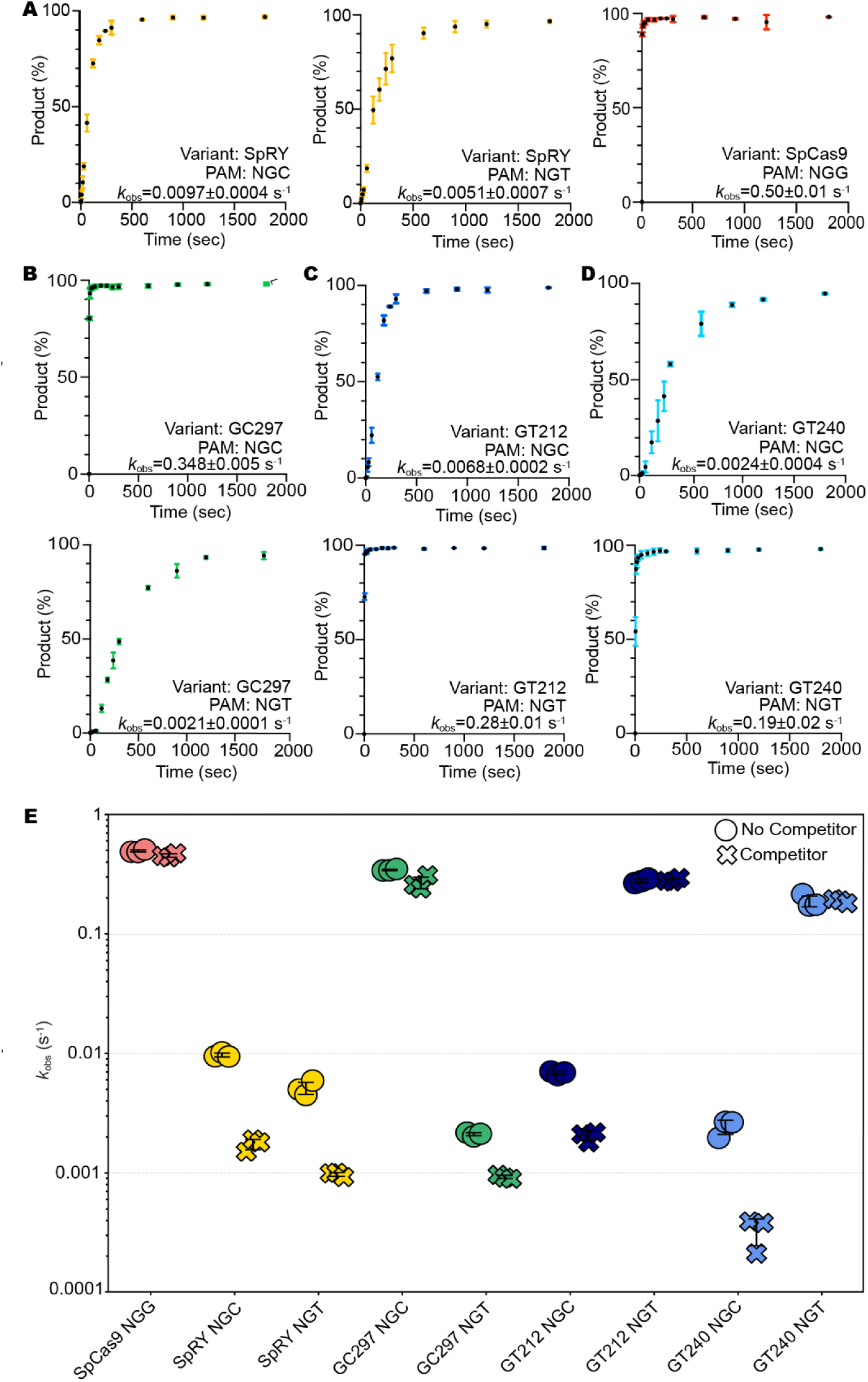
Kinetic analysis of dsDNA cleavage by engineered PAM-selective SpCas9 variants. **(A)** *In vitro* dsDNA cleavage kinetics of SpRY on NGC and NGT PAM substrates and WT SpCas9 on an NGG PAM substrate as controls. **(B–D)** *In vitro* dsDNA cleavage kinetics of engineered PAM-selective variants GC297 **(B)**, GT240 **(C)**, and GT212 **(D)** on NGC and NGT PAM substrates. Cas9 (30 nM active fraction) was incubated with sgRNA and 10 mM Mg^2+^, and reactions were initiated with 10 nM of a 55 bp FAM-labeled dsDNA substrate containing the target protospacer and PAM of interest at 37°C. Aliquots were collected over a 0 s to 2 h time course, where 0 s represents substrate only, and resolved on 15% denaturing urea-PAGE gels (Figure S5B). Data represent three independent replicates; error bars indicate s.e.m. Time courses for the first 30 min are shown. Variant identity, targeted PAM, and observed rate constants (*k*_obs_ ± s.e.m.) from first-order exponential fits are indicated in the bottom right of each graph. **(E)** Summary of *k*_obs_ values for each Cas9 variant and PAM tested in (A–D) without competitor and from parallel reactions performed in the presence of a 1× molar excess of 2.2 kb plasmid competitor DNA (Figure S5C-D). Circles denote *k*_obs_ without competitor; crosses denote *k*_obs_ with competitor.

PAM-relaxed Cas9 variants, such as SpRY, exhibit increased target search times due to a higher frequency of accessible PAM sites, resulting in slower overall cleavage rates. This kinetic bottleneck is particularly pronounced in the presence of excess non-target DNA — a condition more closely reflecting the nuclear environment^15^. To determine whether our top variants exhibit this limitation, we performed in vitro cleavage assays in the presence of 10 nM supercoiled competitor plasmid lacking sgRNA complementarity (Figure 5E and S5C-D). The competitor had minimal impact on wild-type SpCas9, reducing its cleavage rate by just 1.1-fold. SpRY, in contrast, showed a much larger drop in activity, with cleavage rates reduced 5.7-fold on NGC PAMs and 5.3-fold on NGT PAMs. GC297, GT240, and GT212 showed this same resistance to competition, but only at their on-target PAM. GC297’s cleavage rate on its target NGC PAM dropped by just 1.3-fold, while GT212 and GT240 showed no measurable reduction within error on their respective target PAMs. These results suggest that re-specifying PAM preference can largely restore the competitive cleavage kinetics of wild-type SpCas9, but on alternative PAMs.

### Mutational enrichment across the PAM-interacting domain

In addition to identifying top-performing NGC/NGT and NGT/NGC variants, our high-throughput yeast selection platform generated quantitative profiling data for hundreds of thousands of Cas9 variants, enabling mutation-level enrichment quantification by comparing mutation frequencies in top selective variants to those in the full library. Although not exhaustively sampling every substitution, variants carried an average of ∼8 mutations each, providing broad combinatorial coverage of the PAM-interacting domain. Top-performing variants were enriched for mutations at several positions around the PAM-binding interface across mutational backgrounds (Figure S6A and S6B).

## Discussion

CRISPR–Cas9 genome editing is constrained by PAM requirements, limiting access to many therapeutically relevant loci. Although PAM-relaxed variants have expanded the targetable genome, these gains are accompanied by reduced on-target activity, slower on-target cleavage kinetics, and increased off-target editing, suggesting that stringent PAM recognition is intrinsically linked to Cas9 specificity and catalytic efficiency. Here, we demonstrate that re-specifying SpCas9 from NGG toward alternative dinucleotide PAMs can overcome these limitations, expanding targetable sequence space while preserving desirable enzymatic properties. Such programmable PAM re-specification would be valuable for allele-specific editing, base editing, prime editing, and other applications in which access to a precise genomic position depends on the identity of a nearby PAM.

In dominant genetic disorders, a central therapeutic objective is allele-specific disruption of the pathogenic allele while preserving the wild-type allele. However, such strategies are limited by the rarity of disease-linked SNPs that create a PAM difference between alleles that existing Cas9 variants can specifically discriminate between. Using a clinically relevant Huntington’s disease gene (*HTT*) SNP as a proof-of-concept target, we engineered variants with either NGC or NGT PAM specificity. By enabling Cas9 to specifically recognize the noncanonical PAM on the disease allele, these NGC- and NGT-selective variants provide a potential route for allele-specific disruption of *HTT*, and represent a step toward gene editing for a significant percentage of *HTT* patients.

A key component of this study is the development of a yeast selection platform capable of simultaneously screening hundreds of thousands of Cas9 variants for activity on multiple PAMs or protospacer sequences. This high-throughput platform and diversified library allowed us to engineer variants with re- and de-targeted PAM specificities. Top variants were assessed for both on-target and de-targeted PAM activity, directly demonstrating single-nucleotide PAM discrimination. It is likely that additional rounds of screening and generative AI-based diversification could further improve activity, as was recently shown for biochemical training data and PAM engineering^23,53,54^. However, it is notable that a single round of selection yielded variants nearly at biochemical parity with wild-type SpCas9, suggesting past works have identified a large fraction of the critical positions involved in PAM specificity^8,14,18^.

Comprehensive dinucleotide PAM profiling showed selection for activity on one desired PAM, paired with de-selection against a closely related PAM, resulted in concomitant loss of activity across the other 14 dinucleotide PAMs. The broad loss of activity across untargeted PAMs underscores the intrinsic coupling between PAM discrimination and catalytic efficiency whereby enhanced activity toward one substrate comes at the cost of reduced activity toward alternatives^55^. In HEK293Ts, the leading variants retained the PAM preferences observed in yeast, with the GC297 and GC257 Cas9 variants preferentially editing NGC sites and the GT240 and GT212 Cas9 variants preferentially editing NGT sites when tested with distinct guide RNAs targeting multiple endogenous loci. Their increased activity at the selected PAM, reduced activity at the de-targeted PAM, and improved off-target profiles relative to SpRY support the broader principle that therapeutically useful target expansion may be achieved through PAM re-specification rather than PAM relaxation. The variants’ reduced activity across all non-target dinucleotide PAMs, together with their reproducible activity across multiple genomic loci, further extends their broader utility as allele-specific editing tools beyond the specific site and PAM pairing they had originally evolved against.

The biochemical data show that the leading re-specified variants achieved rapid cleavage on their selected PAMs, far outperforming SpRY, with on-target rates approaching those of wild-type SpCas9 on NGG PAMs. They also displayed strong kinetic discrimination between target and non-target PAMs and were less impaired than SpRY by competitor DNA, suggesting that narrowed PAM recognition improves productive substrate engagement under conditions that more closely approximate the complexity of the nuclear environment. Thus, PAM re-specification is not simply a binding-specificity problem, but an integrated enzymological optimization in which PAM recognition, R-loop formation, catalytic efficiency, and off-target suppression are coupled.

More broadly, this work establishes a general framework for engineering novel, stringent PAM specificities beyond those of wild-type SpCas9. This is further highlighted by additional variants that, in our yeast selection system, showed increased activity over SpRY on diverse, difficult-to-target pyrimidine-rich PAMs. A key area for future investigation will be determining whether these variants retain the same robust activity and specificity upon transfer to mammalian cells as the NGC- and NGT-targeting variants analyzed in this study. Together, these findings show that stringent PAM recognition can be reprogrammed, offering a path toward Cas9 editors that combine expanded access with the specificity and catalytic performance required for precision genome editing.

### Limitations

Several limitations remain to be addressed. Although NGC and NGT PAM-targeting variants activity and specificity were characterized across multiple endogenous loci, variant performance may differ across protospacer sequences, chromatin contexts, cell types, and delivery formats not examined here. Cellular off-target analysis, together with *in vitro* genome-wide profiling, supports improved specificity relative to SpRY, but a more complete assessment in disease-relevant cells and under therapeutically relevant expression or delivery conditions will be required. The biochemical assays define intrinsic cleavage properties *in vitro*, but may not fully capture target search, chromatin engagement, or repair outcomes in mammalian nuclei. Similarly, although the mutational profiling and reversion analyses identify residues contributing to NGC/NGT discrimination, the combinatorial nature of the libraries limits definitive assignment of all mutational effects. Although the NGC- and NGT-selective variants have been validated at endogenous mammalian loci, their evaluation at disease-relevant *HTT* alleles is part of an ongoing collaboration. Additional noncanonical PAM-selective variants identified and functionally validated in the yeast screen have not yet been evaluated in mammalian cells, with their evaluation in mammalian systems being part of a separate collaboration. These studies will be important next steps in establishing the therapeutic scope of PAM-re-specified Cas9 enzymes.

## Supplemental figures

**Figure S1.**
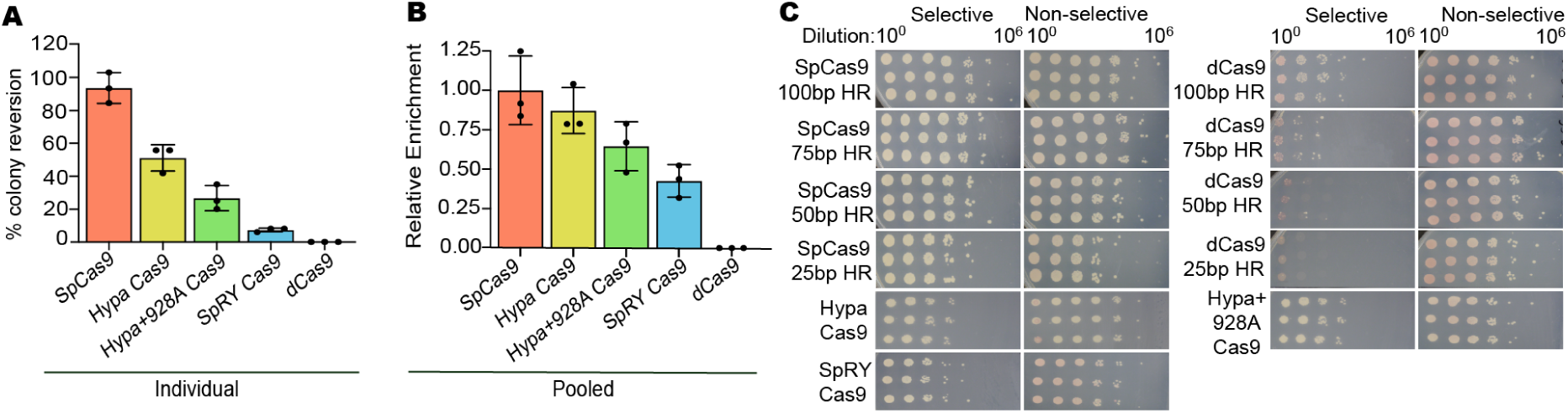
Individual validation of top NGC- and NGT-selective Cas9 variants by yeast cleavage assay. **(A)** Screening of previously published enzymes with known differential NGG PAM activity, including WT SpCas9, HypaCas9, Hypa+928A Cas9, SPRY Cas9, and dCas9. Variants were pooled, transformed into *ADE2* reporter strains with NGG PAM and 25-bp homology arms. Enrichment was computed from barcode abundances in selective relative to nonselective conditions. Enrichment values (± s.e.m.) are from n = 3 independent selections. **(B)** Side-by-side comparison of percent colony reversion for previously published enzymes (WT SpCas9, HypaCas9, Hypa+928A Cas9, SPRY Cas9, dCas9) in the *ADE2* reporter strains with NGG PAM and 25-bp homology arms. Colony reversion efficiency was determined from colony counts on selective (–adenine) versus non-selective (+adenine) plates. Data represent mean ± s.e.m. (n = 3 technical plating replicates, 8 h post-induction). **(C)** Representative titer plates of yeast expressing each variant for 8 h, serially diluted from 10^7^ to 10^0^ CFUs and plated on selective (–adenine) and non-selective (+adenine) media, corresponding to the data quantified in Figure 1C and Figure extended 1A–B. Technical pipetting replicates are shown for each variant/PAM combination.

**Figure S2.**
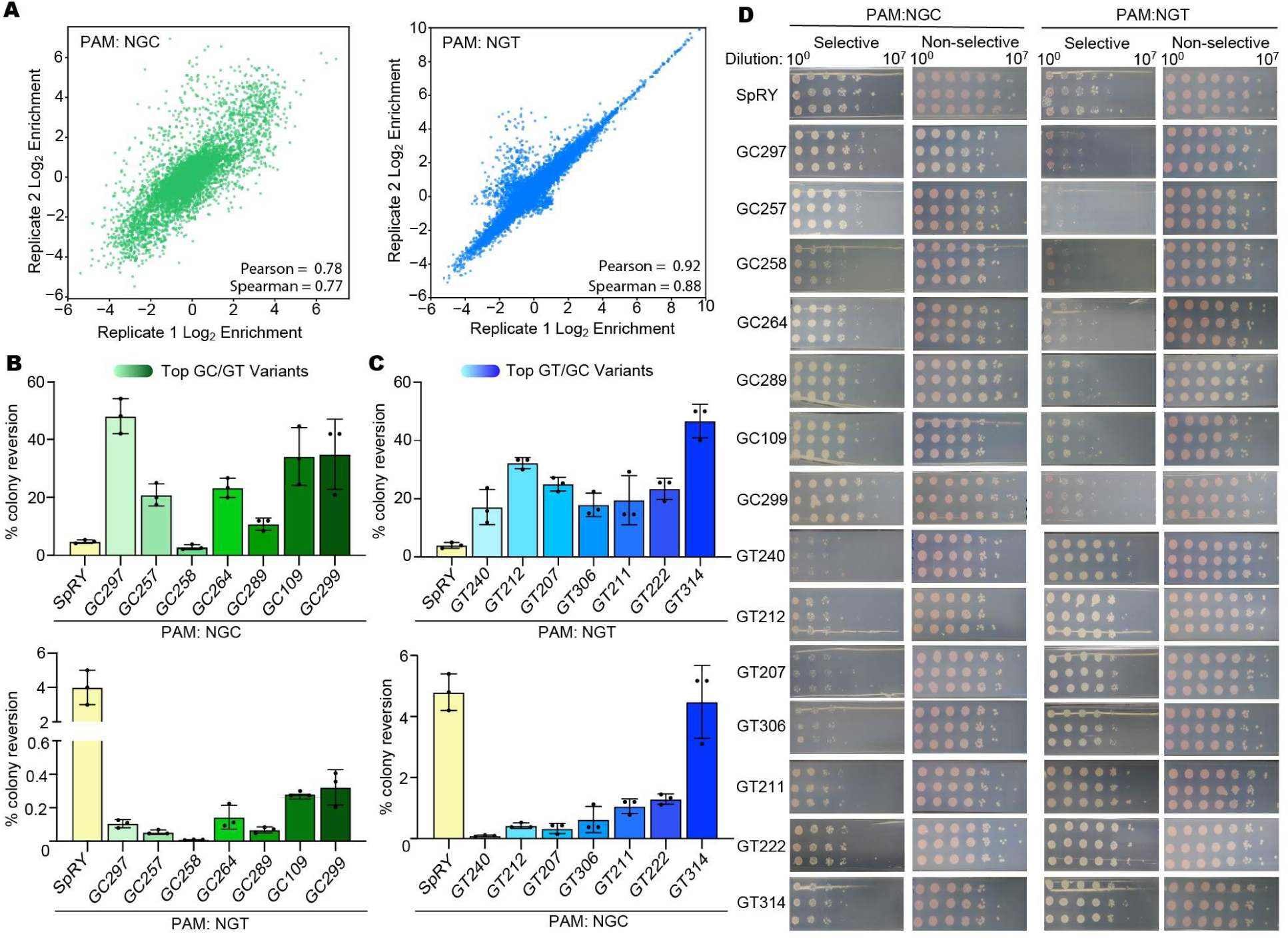
Individual validation of top NGC- and NGT-selective Cas9 variants by yeast cleavage assay. **(A)** Replicate correlation of Log_2_ enrichment scores for the NGC (left) and NGT (right) PAM reporter strain selections shown in Figure 2B, plotting Log_2_ enrichment values from replicate 1 against replicate 2 for all variants in the library. **(B)** Activity of top NGC-selective variants on NGC PAMs, compared to the SpRY control, assessed by percent colony reversion in the yeast cleavage assay. Data represent mean ± s.e.m. (n = 3 technical plating replicates, 8 h post-induction). **(C)** Activity of top NGT-selective variants on NGT PAMs, compared to the SpRY control, assessed by percent colony reversion in the yeast cleavage assay. Data represent mean ± s.e.m. (n = 3 technical plating replicates, 8 h post-induction). **(D)** Representative titer plates of yeast expressing each variant for 8 h, serially diluted from 10^7^ to 10^0^ CFUs and plated on selective (-adenine) and non-selective (+adenine) media. Technical pipetting replicates are shown for each variant/PAM combination.

**Figure S3.**
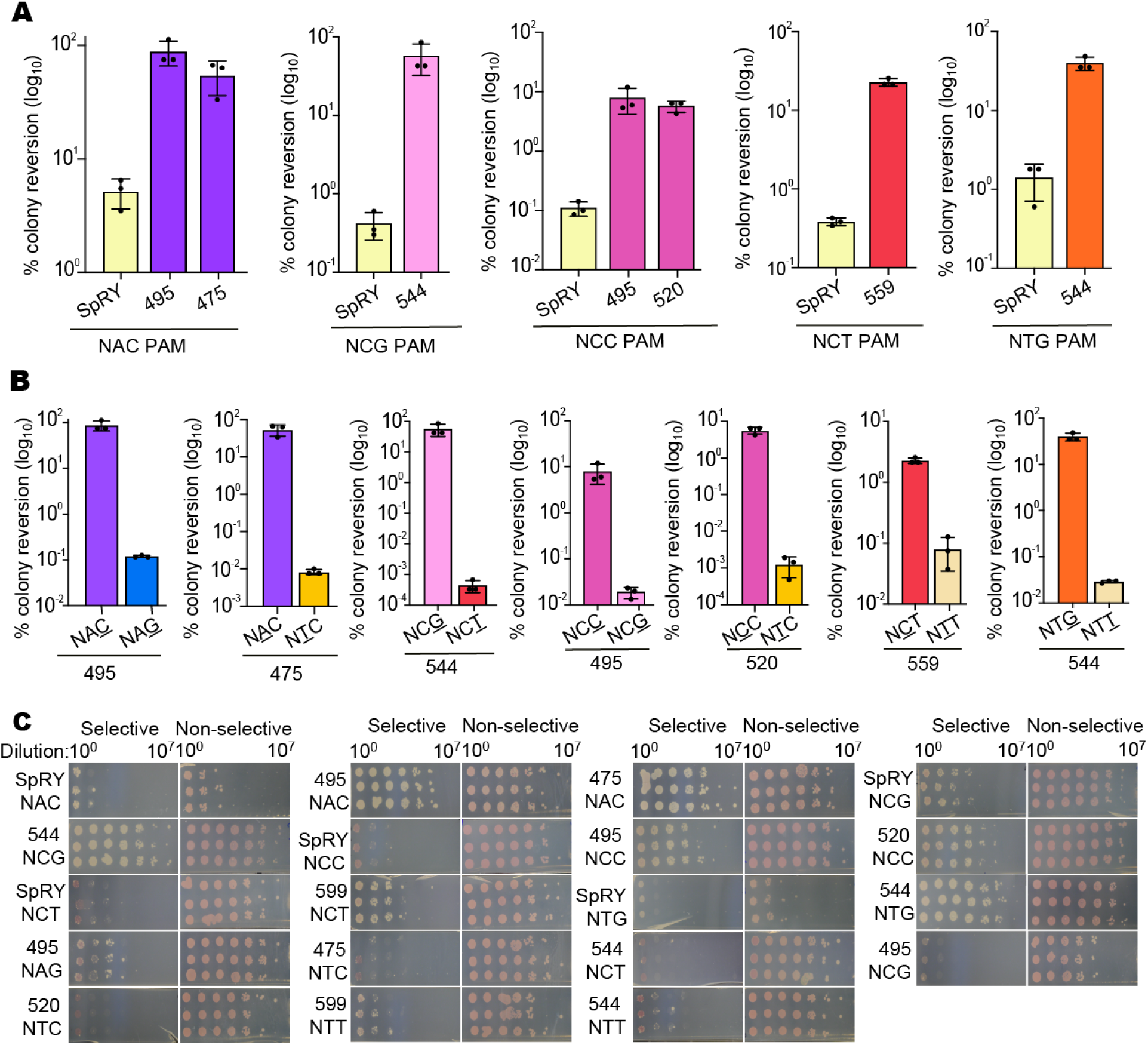
Validation of engineered variants for re-specified recognition of noncanonical dinucleotide PAMs in yeast. **(A)** Percent editing of top-performing variants on their target noncanonical PAMs (NAC, NCG, NCC, NTG, and NCT), compared to the SpRY control, assessed by percent colony reversion in the yeast cleavage assay. Data represent mean ± s.e.m. (n = 3 technical plating replicates, 8 h post-induction). **(B)** Percent editing of each variant on its target PAM compared to the single-nucleotide-differing de-targeted PAM, assessed by percent colony reversion in the yeast cleavage assay. Data represent mean ± s.e.m. (n = 3 technical plating replicates, 8 h post-induction). SpRY is shown in yellow, while all other variants are colored according to the PAM dinucleotide sequence they are being evaluated at, with matching colors representing the same PAM dinucleotide. **(C)** Representative titer plates of yeast expressing each variant for 8 h, serially diluted from 10^7^ to 10^0^ CFUs and plated on selective (-adenine) and non-selective (+adenine) media. Technical pipetting replicates are shown for each variant/PAM combination.

**Figure S4.**
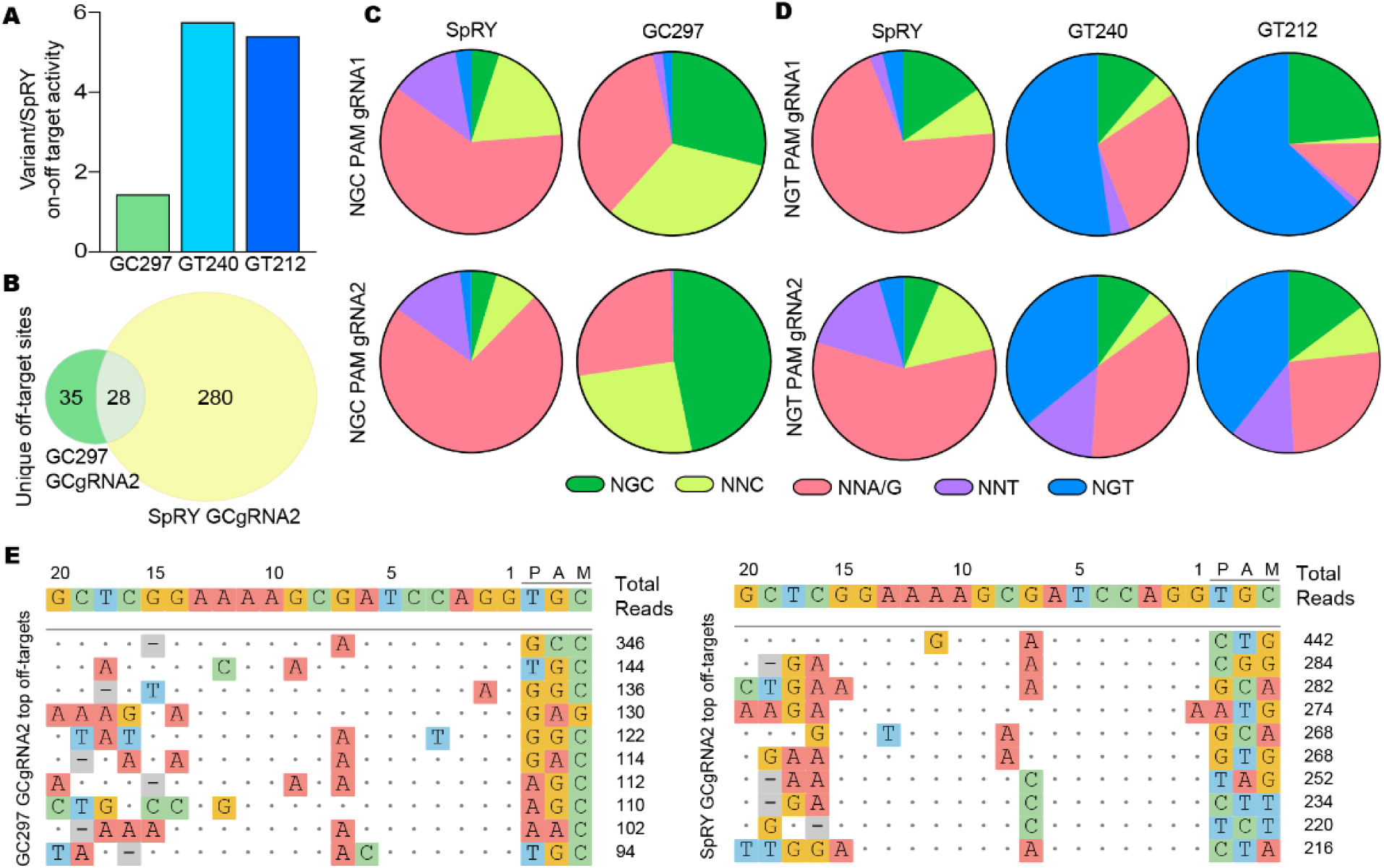
Engineered variants show improved on-target specificity and altered genome-wide PAM preferences relative to SpRY. **(A)** Fold improvement in on-target to off-target editing ratios for the top-performing guide RNA per variant (GC297, GT240, GT212) relative to SpRY. **(B)** Venn diagram representations of CHANGE-seq-detected off-target sites shared between or unique to SpRY and GC297 for GCgRNA2, the variant and gRNA combination showing the smallest improvement in on-target to off-target ratio relative to SpRY. **(C)** Pie charts showing CHANGE-seq-detected genome-wide off-target sites by PAM class for SpRY and GC297, each paired with two sgRNAs targeting genomic loci containing NGC PAMs in HEK293T cells. **(D)** Pie charts showing CHANGE-seq-detected genome-wide off-target sites by PAM class for SpRY, GT212, and GT240, each paired with two sgRNAs targeting genomic loci containing NGT PAMs in HEK293T cells. **(E)** Representative top off-target sites identified by CHANGE-seq for SpRY and GC297 paired with GCgRNA2. The top 10 off-target sites and their associated PAMs are shown, demonstrating stronger enrichment for NGC PAMs and, to an even greater extent, NNC PAMs among the highest-ranked off-target sites compared with the overall off-target population. CHANGE-seq was performed with n = 2 replicates for all variants and sgRNAs, and data shown reflect both replicates.

**Figure S5.**
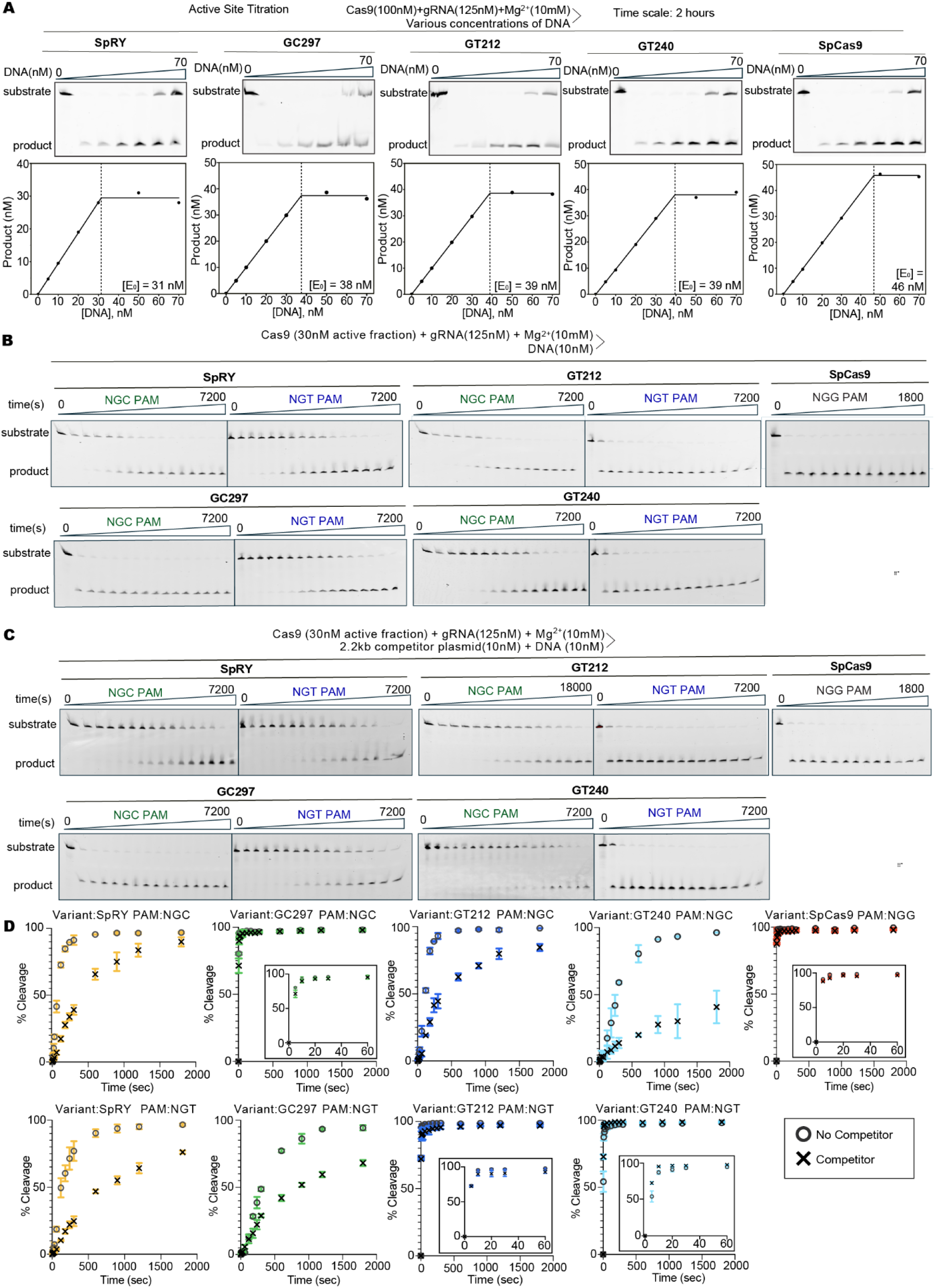
Determination of active enzyme fraction and *in vitro* dsDNA cleavage kinetics for engineered PAM-targeting variants. **(A)** Active site titration was performed by incubating Cas9 (100 nM) with gRNA (125 nM) and 10 mM Mg^2+^, with variable concentrations (0–70 nM) of a FAM-labeled dsDNA substrate containing the complementary protospacer and PAM of interest, at 37°C. Reactions were allowed to proceed for 2 h to reach completion, and cleavage products were resolved on 15% denaturing urea-PAGE gels. The concentration of active enzyme ([E₀]) was determined from the inflection point of the resulting product versus DNA plot, indicated by the vertical dashed line. **(B)** Cas9 (30 nM active fraction) was incubated with gRNA (125 nM) and 10 mM Mg^2+^, then reactions were initiated with 10 nM 55 bp FAM-labeled DNA substrate containing the complementary protospacer and PAM of interest, at 37°C. Time-course fractions were collected at 0 s, 5 s, 10 s, 20 s, 30 s, 1 min, 2 min, 3 min, 4 min, 5 min, 10 min, 15 min, 20 min, and 30 min (“0” represents substrate only) and resolved on 15% denaturing urea-PAGE gels; a representative gel from three independent replicates is shown. Additional 1 h and 2 h timepoints were collected for all variants except WT SpCas9. Gels for reactions without competitor are shown for SpRY, GC297, GT212, and GT240 on NGC and NGT PAMs, and for WT SpCas9 on its native NGG PAM. **(C)** Gels for reactions performed in the presence of a 1× 2.2 kb competitor plasmid, shown for the same variants and PAMs as in (B); for GT240 on the NGC PAM with competitor, additional timepoints at 3, 4, and 5 h are included to capture its slower cleavage kinetics. A representative gel from three independent replicates is shown. **(D)** *In vitro* dsDNA cleavage kinetics corresponding to the competitor gels in (C) for control variants SpRY on NGC and NGT PAMs, and SpCas9 on its native NGG PAM and GC297, GT212 and GT240 on NGC and NGT PAMs. The first 30 minutes of cleavage is shown. Crosses denote reactions performed with competitor DNA; circles denote reactions performed without competitor DNA and are shown for comparison. Zoomed-in plots of the first 300 seconds of cleavage are shown for WT SpCas9 and engineered variants on their corresponding on-target PAM substrates. Data represent three independent replicates; error bars indicate s.e.m.

**Figure S6.**
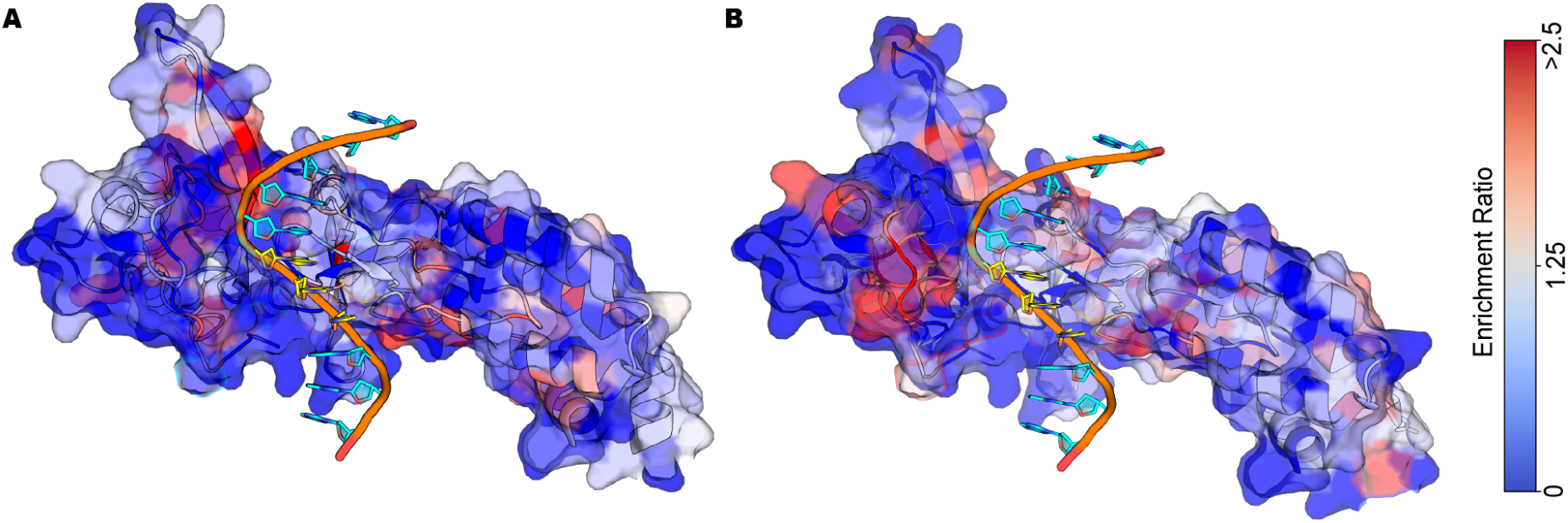
PAM-selective mutational enrichment across the Cas9 PAM-interacting domain. Mutational enrichment of top PAM-selective variants mapped onto the Cas9 PAM-interacting domain (PID). Enrichment ratios of amino acids differing from those present in SpRY at each position in **(A)** the top 1% of NGC/NGT-selective and **(B)** top 1% of NGT/NGC-selective variants relative to the full library were projected onto the Cas9 PID structure (PDB 4UN3). Color scale indicates enrichment score, with red representing positions with higher enrichment values, as shown in the scale bar. DNA is shown in orange, with PAM bases highlighted in yellow.

## Methods

### Plasmid construction

The plasmid p415-GalL-Cas9-CYC1t^56^ (Addgene #43804) was used as the backbone for yeast Cas9 expression. Catalytically inactive dCas9 (D10A/H840A), HYPA Cas9 (N692A/M694A/Q695A/H698A), and HYPA+R928A variants were generated by introducing point mutations via blunt-end ligation. SpRY carrying the high-fidelity mutations N497A, R661A, Q695A, and Q926A was cloned into p415-GalL-Cas9-CYC1t in place of wild-type Cas9 by PCR and Gibson assembly using Q5 High-Fidelity Polymerase (NEB). All constructs were verified by Sanger sequencing at the UC Berkeley DNA Sequencing Facility (Barker Hall). sequencing.

Rationally designed Cas9 libraries were constructed from an oligonucleotide pool (Twist Bioscience) covering the PAM-interacting domain (PID) of Cas9. Residues 1333 and 1335, which contact the second and third PAM positions respectively, were biased toward amino acids known from the zinc finger, TALEN, and Cas9 literature to make base-specific contacts^8,39–46^. Positions known to influence PAM recognition (1135, 1136, 1218, 1219, and 1337) were randomized to NNN to sample the full amino acid space at high mutational frequency. Random mutations were subsequently introduced across the PID using the GeneMorph II Random Mutagenesis Kit (Agilent Technologies) with 5 ng template per 50 μL reaction over 30 cycles, achieving a final average of approximately 8 mutations per PID variant, inclusive of mutations introduced by the Twist oligonucleotide pool. A 20-bp barcode was incorporated into the reverse primer, providing each unique Cas9 variant with a distinct barcode at the Cas9 C-terminus.

The mutagenized PID was cloned into p415-GalL-Cas9-CYC1t^56^ (Addgene #43804) by Golden Gate assembly using flanking BsaI sites, replacing a GFP stuffer fragment. Endogenous BsaI sites within the backbone had been removed by a multi-fragment Gibson assembly performed simultaneously across all sites. Assemblies were transformed into TOP10 electrocompetent *E. coli*, which was selected for its higher transformation efficiency to maximize library diversity, yielding a primary library of ∼10⁷ CFUs as estimated from titer plates. The library was plated on LB-carbenicillin plates at a density permitting individual colony resolution and colonies were counted using the Analyze Particles function in Fiji^57^ (size: 120–∞ pixels; circularity: 0.35–1.00) following binary thresholding and watershed segmentation, to bottleneck the library to a final size of ∼2×10⁵. Plasmid DNA was purified from the bottlenecked library. Variants were mapped to their respective barcodes by long-read sequencing (PacBio Sequel II) by QB3 at UC Berkeley. All reads were aligned to a reference plasmid and barcode sequences extracted using Minimap2^58^ (version 2.26) and SAMtools^59^ (version 1.19). Subalignments were made for all reads with a given barcode and a consensus sequence was created using SAMtools for all barcodes with at least two reads for PacBio sequencing. Barcodes not 20bp in length were discarded, for a total of 106675 variants mapped.

The expression cassette for the sgRNA was obtained from p426-SNR52p-gRNA.CAN1.Y-SUP4t^56^ (Addgene #43803); the original spacer sequence was replaced with the desired target sequence by PCR and Gibson assembly using Q5 High-Fidelity Polymerase (NEB). Constructs were transformed into Turbo competent *E. coli* (NEB) and verified by Sanger sequencing.

The mammalian Cas9 expression vector was assembled by Gibson assembly using Q5 High-Fidelity Polymerase (NEB). The Cas9 coding sequence excluding the PAM-interacting domain, a GFP placeholder fragment flanked by BsaI sites in place of the PAM-interacting domain, and the mCherry sequence were cloned into a backbone plasmid provided by Cynthia Terrace of the Savage lab (University of California, Berkeley). The GFP placeholder fragment flanked by BsaI sites was in place of the PID allows efficient Golden Gate cloning of PID variants from top library hits. The sgRNA mammalian expression vector pJRH051^60^ (Addgene #171625) contained flanking BsmBI sites, which permitted us to clone in all our spacer sequences. Spacers were annealed from oligonucleotides and cloned via Golden Gate cloning (IDT). Constructs were transformed into Turbo competent *E. coli* (NEB) and verified by Sanger sequencing.

Bacterial expression vectors for top variants were generated by assembling gBlocks (IDT) encoding the bacterially codon-optimized PID containing the desired mutations into the SpRY bacterial expression vector pSHS207^48^ (Addgene #101199) by Golden Gate assembly using Q5 High-Fidelity Polymerase (NEB). Constructs were transformed into Turbo competent *E. coli* (NEB) and verified by Sanger sequencing.

To assess the contribution of individual mutations in top-performing variants identified from the yeast selection screen (GC297 and GT240), single reversion mutations to the SpRY parental sequence and substitutions of interest identified from mutational enrichment analysis were introduced into the yeast expression vector by Golden Gate assembly using flanking BsaI sites. Constructs were transformed into Turbo competent *E. coli* (NEB) and verified by Sanger sequencing prior to transformation into reporter yeast strains as described above.

### Reporter yeast strain creation

Yeast reporter strains were generated using the *delitto perfetto* approach to edit the *ADE2* genomic locus according to published protocols^35^. An intermediate *ADE2* knockout was derived from *Saccharomyces cerevisiae* BY4741 (ATCC 201388) using the CORE cassette GSKU, excluding Gal-I-SceI. The intermediate strain was then co-transformed with linearized DNA containing the *ADE2* coding sequence split by insertion of the desired spacer, PAM sequence, and a stop codon, flanked by duplicated homology regions of 100 bp, 75bp, 50 bp, or 25 bp. Following this initial comparison, the 25 bp homology region was selected for all subsequent experiments as it yielded the greatest dynamic range between active and catalytically inactive Cas9, owing to reduced background activity from the inactive control. A plasmid carrying constitutively expressed SpyCas9 targeting the CORE cassette was co-transformed to induce DSBs and drive repair from the linear DNA template. Successful integration was confirmed by PCR and Sanger sequencing, after which strains were cured of the Cas9 plasmid. A unique barcode was incorporated at the *ADE2* C-terminus to identify each strain. Using this approach, we generated 18 *ADE2* reporter strains: all 16 possible dinucleotide combinations at the second and third PAM positions (NN PAM series), plus two additional strains harboring NGC and NGT PAMs paired with a spacer targeting an allele-specific exonic SNP in the *HTT* gene relevant to allele-specific therapeutic silencing (NGC_HD and NGT_HD).

### Selection assays in reporter yeast strain

sgRNA plasmids (carrying a *URA3* marker) were transformed into *Saccharomyces cerevisiae* BY4741 *ADE2* reporter strains using 2 μg plasmid via the lithium acetate/single-stranded carrier DNA/PEG method^61^ and plated on −ura solid medium to select for stable transformants. For evaluation of individual Cas9 variants, 5 μg of Cas9 plasmid (carrying a *LEU2* marker) was transformed by the same method.

For Cas9 libraries, the Cas9 plasmid was linearized by SapI digestion and the Cas9-containing fragment was gel-extracted. A PCR product containing the remaining vector sequence with 60 bp homology arms to the Cas9 fragment was generated. Yeast were grown to OD_600_ 1.5–1.6, pelleted, and washed twice with ice-cold water and once with ice-cold electroporation buffer (1 M sorbitol, 1 mM CaCl₂). Cells were conditioned in 20 mL conditioning solution (0.1 M LiAc, 10 mM DTT) at 30°C for 30 min with shaking, then washed and resuspended in electroporation buffer. 5 μg linearized Cas9 fragment and 1 μg PCR product were mixed with 400 μL electrocompetent cells, incubated on ice for 5 min, and electroporated at 2.5 kV, 25 μF in a 0.2 cm cuvette (Bio-Rad GenePulser) alongside SpRY and catalytically inactive Cas9 controls. Cells were recovered in 8 mL YPD/1 M sorbitol (1:1) at 30°C for 1 h with shaking, then resuspended in liquid SCD −leucine −uracil medium. Library expression vectors were assembled *in vivo* by gap repair homologous recombination.

All transformants — both library electroporations and individual lithium acetate/single-stranded carrier DNA/PEG transformations — were recovered overnight at 30°C in liquid SCD −leucine −uracil medium to select for both plasmids. The following morning, cultures were washed three times in galactose medium to remove residual glucose, then induced in liquid SCD −leucine −uracil medium supplemented with 2% (w/v) galactose. An initial experiment determined 8 hours to be optimal for dynamic range; all subsequent inductions were carried out for 8 h at 30°C. Cultures were supplemented with adenine (200ug/ml) prior to plating on selective (−adenine −leucine −uracil) and non-selective (+adenine −leucine −uracil) SCD solid medium on bioassay dishes (Thermo Fisher). Plates were incubated at 30°C for 48 h, after which colonies were scraped. Colony PCR was performed to amplify the barcoded region of the Cas9 plasmid by first heating yeast in 25 mM NaOH at 98°C for 15 min, centrifuging briefly, and using the clarified lysate as template for Q5 High-Fidelity DNA Polymerase (NEB). PCR products were purified with AMPure XP beads (Beckman Coulter). Illumina sequencing (NovaSeq, paired-end 150 bp) was performed by Novogene (Davis, CA) at an average depth of ∼60million read/sample. Barcode frequencies were quantified using a custom Python script. Briefly, 20-nt barcode sequences were extracted by regular expression matching, counted, normalized to sequencing depth, and expressed as enrichment values calculated as the log_2_ ratio of reads in selective versus non-selective conditions. Barcodes were filtered out if none or one of the nonselective replicates met the >3-read threshold, and further filtered out as outliers if replicate standard deviation exceeded the mean. For replicate correlation analyses, low-count data were filtered out prior to plotting, retaining only barcodes with a minimum raw read count of 25. Results were plotted with matplotlib^62^.

To characterize top variants from the screen, variants were cloned from the library using their unique barcode as a reverse primer and re-cloned into the yeast expression vector. Variants were transformed into reporter strains carrying the desired ON- or OFF-target PAM using 5 μg Cas9 plasmid via the lithium acetate/single-stranded carrier DNA/PEG method^61^ and induced as above. Following the 8 h induction, 10-fold serial dilutions were prepared in triplicate and plated on selective (−adenine −leucine −uracil) and non-selective (+adenine −leucine −uracil) SCD solid medium. After 48 h at 30°C, CFUs were counted and colony reversion was calculated as (CFUs on selective / CFUs on non-selective) × 100. Data were plotted using GraphPad Prism (v10.6.1).

To obtain complete dinucleotide PAM activity profiles, 16 *ADE2* reporter strains — each harboring one of the 16 possible dinucleotide PAMs at the second and third PAM positions and a unique barcode at the *ADE2* C-terminus — were pooled. Each variant of interest was transformed into the pooled 16-strain library via the lithium acetate/single-stranded carrier DNA/PEG method^61^. Selection, colony scraping, and yeast PCR’s were performed as described for the library screen, except that the barcode at the *ADE2* genomic locus C-terminus was amplified. Amplicon sequencing was performed at the MGH Sequencing Core. For each PAM, enrichment was calculated as the fraction of reads corresponding to that PAM in the selective condition divided by the corresponding fraction in the non-selective condition. The resulting values were then normalized to sum to 1, such that each value represented the fractional contribution of that PAM to the total enrichment

### Mammalian genome editing

HEK293T cells (UC Berkeley Cell Culture Facility) were cultured in DMEM (Corning) supplemented with 10% fetal bovine serum (Gibco). Cells were seeded at ∼20,000 cells/well in 96-well plates 16–24 h before transfection. Transfection mixes were prepared by combining 80 ng Cas9 mammalian expression vector and 20 ng sgRNA mammalian expression vector with 9 μL Opti-MEM I (Thermo Fisher) and 0.3 μL TransIT-293, incubated at room temperature for 30 min, and added dropwise to cells.

Five days post-transfection, cells were harvested and lysed in QuickExtract DNA Extraction Solution (Biosearch Technologies) by sequential incubation at 65°C for 20 min and 98°C for 20 min. Genomic loci were amplified from lysates by two rounds of PCR using Q5 High-Fidelity Polymerase (NEB): the first to amplify target loci and attach adapter sequences, and the second to add Illumina index and P5/P7 sequences. PCR products were purified with AMPure XP beads (Beckman Coulter), pooled at equimolar concentrations, and sequenced on an Illumina NextSeq or MiSeq (150-bp paired-end) at the Innovative Genomics Institute (IGI) NGS Core. Editing frequencies were quantified using CRISPResso2 (v2.3.1)^63^. Editing data were plotted using GraphPad Prism (v10.6.1).

For off-target analysis, Cas-OFFinder (v2.4.1; http://www.rgenome.net/cas-offinder/) ^64^ was used to predict potential genomic off-target sites. These predicted sites were amplified and sequenced as described above to compare the off-target activity of our NGC- and NGT-targeting variants against SpRY. Editing data were plotted using GraphPad Prism (v10.6.1).

### Protein expression and purification

Cas9 variants were expressed and purified following a modified protocol based on Cofsky et al^65^. *E. coli* Rosetta (DE3) cells (Sigma-Aldrich) harboring bacterial expression plasmids encoding each Cas9 variant were grown in Terrific Broth (TB; ThermoFisher Scientific) supplemented with ampicillin (0.1 mg/mL) at 37°C, diluted 1:200 from overnight starter cultures. At OD₆₀₀ = 0.6, expression was induced with 0.5 mM IPTG overnight at 16°C. Cells were pelleted, resuspended in lysis buffer (20 mM HEPES pH 7.5, 500 mM KCl, 10 mM imidazole, 10% glycerol, 1 mM TCEP, cOmplete EDTA-free protease inhibitor cocktail (Roche)), and lysed by sonication. Lysates were clarified by centrifugation at 18,000 x g for 30 minutes, and the resulting supernatant was applied to Ni-NTA resin (QIAGEN), washed (20 mM HEPES pH 7.5, 500 mM KCl, 30 mM imidazole, 5% glycerol, 1 mM TCEP), and eluted with 300 mM imidazole in the same buffer. His-tags were removed by TEV protease cleavage during overnight dialysis at 4°C (20 mM HEPES pH 7.5, 300 mM KCl, 30 mM imidazole, 5% glycerol, 1 mM TCEP). Protein was further purified over a HiTrap Heparin HP column (Cytiva) using a 300 mM–1 M KCl gradient, then aliquoted, snap-frozen, and stored at −80°C. Purification was assessed by SDS-PAGE on 4–20% Criterion TGX Precast Gels (Bio-Rad) with Thermo Scientific Spectra Multicolor Broad Range Protein Ladder(ThermoFisher Scientific), stained with InstantBlue Coomassie (Abcam) and imaged on a ChemiDoc system (Bio-Rad). Purified wild-type SpCas9 protein was purchased from the QB3 Macrolab (University of California, Berkeley)

### CHANGE-seq

To evaluate genome-wide off-target activity, CHANGE-seq^52^ was performed on GC297, GT212, GT240, and SpRY using purified proteins complexed with sgRNAs (IDT) targeting genomic loci adjacent to their respective on-target PAMs. Analyzed targets included an NGC PAM–dependent locus (GCgRNA1) previously characterized with SpRY by GUIDE-seq^14^, the disease-relevant *HTT* locus (GTgRNA2), and an additional high-activity endogenous locus from HEK293T experiments for each PAM. Genomic DNA from NA12878 human cells was extracted (PureGene Cell Kit, Qiagen) and tagmented with Tn5 transposase to ∼400 bp fragments. Gaps were repaired, and DNA was circularized with T4 DNA ligase; linear DNA was removed by exonuclease digestion. Circularized DNA (125 ng) was cleaved *in vitro* with Cas9:sgRNA RNPs (90uM:180uM) A-tailed, and ligated to Illumina adapters. Libraries were barcoded by PCR (NEBNext Multiplex Oligos, Q5 polymerase), purified (Ampure XP), and sequenced as 2×150 bp paired-end reads on a NextSeq 2000 (∼5-8 million reads/sample). Sequence analysis was performed using a modified version of CHANGE-seq analysis software (https://github.com/Interventional-Genomics-Unit/IGU_CHANGEseq). First, FASTQs were processed by quality-filtering (fastp v0.24.0) and trimmed to remove transposon and Illumina adapter sequences using the Cutadapt (v3.4.23) command ‘cutadapt -a CTGTCTCTTATACACATCTACGTAGATGTGTATAAGAGACAG -A CTGTCTCTTATACACATCTACGTAGATGTGTATAAGAGACAG -o [R1_OUT] -p [R2_OUT] -Z --overlap 35 -e 0.15 -m 30 -j 48 [R1] [R2]’ . Trimmed reads were then aligned to the human reference genome GRhg38.p14 using BWA (v0.7.17). Aligned reads were classified as edited in both RNP-treated and untreated control samples according to the following criteria: (1) had mapping quality threshold, MAPQ < 40 (2) outward-facing paired-end alignment had a breakpoint distance ≤ 5 bp (3) aligned reads contained the spacer sequence along with a 5’ “NNN” PAM below an edit distance ≤ 7 with a max bulge threshold ≤1 and the matching sequence was within +/– 30 bp or aligned sequence ends. Lastly, to account for variability between replicates, raw read counts of identified sites were normalized using median normalization. Final sites were reported if at least one replicate contained a site with ≥6 reads.

### Nucleic acid preparation

55-nt DNA duplexes were prepared from HPLC-purified oligonucleotides (Integrated DNA Technologies), using a 20-nt spacer sequence adapted from Hibshman et al.^17^ and NGG, NGC, or NGT PAM sequences. For cleavage assays, 6-FAM-labeled target strands were annealed to unlabeled non-target strands at a 1:1.15 molar ratio in annealing buffer (10 mM Tris-HCl pH 8, 50 mM NaCl, 1 mM EDTA) by heating to 95°C for 5 min, then cooling to room temperature over 70 min. sgRNA was transcribed from an IDT-purchased DNA template using the NEB HiScribe™ T7 High Yield RNA Synthesis Kit (NEB #E2040S/L). Reactions were incubated at 37°C for 4 h. RNA was purified by 15% denaturing urea polyacrylamide gel electrophoresis (urea-PAGE). Gel slices containing RNA were crushed and soaked in DEPC-treated water overnight at 4°C, then washed six times with DEPC water. The plasmid (pGGAselect) used in competition cleavage assay was ordered from New England Biolabs and prepared and gifted by Honglue Shi of the Doudna lab at UC Berkeley^15^.

### DNA cleavage kinetics

All cleavage reactions were performed in 1× cleavage buffer (20 mM Tris-Cl pH 7.5, 100 mM KCl, 5% glycerol, 1 mM DTT) at 37°C.

#### Active fraction determination

To determine the active fraction of each Cas9 purification, an active site titration was performed by pre-incubating 125 nM sgRNA, 5 mM Mg²⁺, and 5–70 nM Cas9 (5, 10, 20, 30, 50, or 70 nM) in 1× cleavage buffer for 5 min, then initiating cleavage by adding the Cas9-gRNA-Mg²⁺ complex to 10 nM 5′-FAM-labeled 55-bp DNA substrate containing a perfectly matched protospacer. After 2 hours, reactions were quenched with an equal volume of 2× quench buffer (94% formamide, 30 mM EDTA, 400 μg/mL heparin, bromophenol blue), heated at 90°C for 5 min, and resolved on a 15% urea-PAGE gel. Gels were scanned on a Typhoon (Amersham/GE Healthcare) with 488 nm excitation and a Cy2 emission filter (525BP20). Band intensities were quantified using Bio-Rad ImageLab 6.1 and data were fitted to a piecewise linear model to determine active fraction concentration as in Gong et al.^66^ (active fraction concentration range: 0.31-0.46).

#### Cleavage kinetics

To measure cleavage kinetics, 125 nM sgRNA, 5 mM Mg²⁺, and 30 nM active Cas9 were pre-incubated in 1× cleavage buffer for 5 min, then cleavage was initiated by addition of 10 nM 5′-FAM-labeled 55-bp DNA substrate containing a perfectly matched protospacer and the PAM of interest. Samples were quenched at defined timepoints, resolved by 15% urea-PAGE, and imaged and quantified as described above. Data were fitted to a first-order exponential model to determine observed cleavage rate constants (*k*_obs_). For competition assays, target dsDNA and the 2.2 kb pGGAselect plasmid competitor were pre-mixed at a 1:1 molar ratio prior to RNP addition.

### Mutational enrichment analysis

To quantify position-specific mutational contributions to NGC and NGT PAM selectivity, per-variant enrichment scores were calculated as described above. Variants were ranked by their NGC/NGT enrichment ratio (NGC enrichment score divided by NGT enrichment score) and vice versa for NGT/NGC selectivity, to determine the top 1% of NGC>NGT and NGT>NGC variants respectively. Per-position mutational enrichment was calculated by dividing the frequency of mutations at each position within the top 1% of NGC>NGT and NGT>NGC variants respectively by the frequency of mutations at that position across the full library. These per-position enrichment values were mapped onto the Cas9 PAM-interacting domain structure (PDB: 4UN3) using PyMOL to visualize spatially enriched residues.

